# Universal method for generating knockout mice in multiple genetic backgrounds using zygote electroporation

**DOI:** 10.1101/2023.04.10.536207

**Authors:** Tomohiro Tamari, Yoshihisa Ikeda, Kento Morimoto, Keiko Kobayashi, Saori Mizuno-Iijima, Shinya Ayabe, Akihiro Kuno, Seiya Mizuno, Atsushi Yoshiki

**Author notes:** Corresponding authors: Shinya Ayabe Experimental Animal Division, RIKEN BioResource Research Center, 3-1-1 Koyadai, Tsukuba, Ibaraki 305-0074, Japan, Akihiro Kuno Department of Anatomy and Embryology, University of Tsukuba, 1-1-1 Tennodai, Tsukuba, Ibaraki 305-8575, Japan, Seiya Mizuno Laboratory Animal Resource Center and Trans-Border Medical Research Center, University of Tsukuba, 1-1-1 Tennodai, Tsukuba, Ibaraki 305-8575, Japan. These authors contributed equally to this work (Tomohiro Tamari, Yoshihisa Ikeda, Kento Morimoto).

## Abstract

Genetically engineered mouse models are essential tools for understanding mammalian gene functions and disease pathogenesis. Genome editing allows for the generation of these models in multiple inbred strains of mice without backcrossing. Zygote electroporation dramatically removed the barrier for introducing the CRISPR-Cas9 complex in terms of cost and labour. However, the editing conditions and protocols to produce knockout lines have been optimised for a limited number of strains or stocks. Here, we demonstrate a novel and universal approach for generating knockout mice in multiple inbred strains. By combining in vitro fertilisation and electroporation, we obtained founders for knockout alleles in 8 common inbred strains. Long-read sequencing analysis detected not only intended mutant alleles but also differences in read frequency of intended and unintended alleles among strains. Successful germline transmission of knockout alleles demonstrated that our novel approach can establish mutant mice targeting the same locus in multiple inbred strains for phenotyping analysis, contributing to reverse genetics and human disease research.

**Summary statement:** Universal method for zygote genome editing in multiple inbred mouse strains allows for generation of novel mutant mice for understanding mammalian gene function and human disease pathogenesis.

## Introduction

An inbred strain of laboratory mice, in which all loci are essentially homozygous, is defined as a strain maintained through 20 or more generations of brother-to-sister mating. Recently, various detailed genome assemblies have been developed, providing information on various inbred mouse strains (Lilue et al., 2018). Each inbred strain exhibits its own unique phenotype and its utilization for the replication of the same or multiple experiments allows the uncovering of the genetic and environmental effects on phenotypes (Li and Auwerx, 2020). The severity and penetrance of abnormal phenotypes can differ when the same target gene is disrupted in different inbred mouse genetic backgrounds (Hide et al., 2002; Widmayer et al., 2020), providing an opportunity to gain insights into the mechanisms of diseases involving multiple genes.

Establishing mouse embryonic stem cells with the potential for germline transmission from various inbred strains has been technically difficult and time-consuming (Tanimoto et al., 2008; Iijima et al., 2010). In addition, embryonic stem cell-derived sequences will remain surrounding the targeted locus even after extensive backcrossing of different host strains. However, zygote genome editing has eliminated these limitations. In particular, genome editing using CRISPR-Cas9 has been adopted in various animal models, including laboratory mice, owing to its high mutation induction efficiency (Wang et al., 2013). The advent of zygote electroporation, such as the TAKE method which does not require advanced techniques, has significantly increased the versatility of genome editing (Kaneko et al., 2014).

Although there have been reports of zygote electroporation using several inbred mouse strains (Nakano et al., 2022), no study has examined whether knockout of the same target gene under the same conditions is possible in multiple mouse strains. We here demonstrate that our method combining in vitro fertilisation and the TAKE method is capable of inducing the knockout of genes in 8 common and easily accessible inbred mouse strains: BALB/cAnNCrlCrlj (BALB/c), NC/NgaTndCrlj (NC), CBA/J (CBA), C3H/HeNCrl (C3H), SJL/J (SJL), DBA/1JNCrlj (DBA1), DBA/2NCrl (DBA2), and C57BL/6NCrl (B6N). We also demonstrate that long-read sequence analysis is effective for checking the frequency of appearance of intended and unintended mutant alleles. Moreover, we confirm that the introduced knockout alleles are consistently inherited by progenies. Schemes that disrupt the same target genes through a universal method in different inbred mouse genetic backgrounds will facilitate the advancement of complex genetic research.

## Results

### In vitro fertilization and embryo transfer in multiple inbred mouse strains

Zygote electroporation has been used to generate mutant rodent strains (Kaneko, 2018). In vitro fertilisation (IVF) following super-ovulation induction reduces the number of females required for embryo collection (Mochida, 2020). Although the response to super-ovulation treatment, IVF, and developmental rates differ in mice depending on their genetic background (Kito et al., 2004; Byers et al., 2006; Ostermeier et al., 2008), the combined effects of methyl-β-cyclodextrin and reduced glutathione on fertilisation and birth rates have been scarcely reported in inbred strains other than C57BL/6 (Takeo and Nakagata, 2011). Therefore, as a preliminary step in genome editing, we examined the efficiency of super-ovulation induction, IVF, and developmental rates. We observed that all strains showed fertilisation rates >80%, ranging from 83.9% (DBA1) to 98.1% (C3H) (Table 1). We further found that the number of fertilised zygotes per female ranged from 8.2 (CBA) to 21.3 (C3H), and 2-cell developmental rate ranging from 97.3% (SJL) to 100% (NC). Interestingly, the achieved pregnancy rate was 100 % for all strains except for B6N (66.7%). We obtained newborns from all tested strains, with birth rates ranging from 16.7% (DBA2) to 70.0% (NC). Pups did not exhibit any phenotypical abnormalities at the age of weaning. These results suggested that our method of combining super-ovulation, IVF, and embryo transfer was effective for conducting zygote genome editing in all 8 inbred mouse strains.

**Table 1.** In vitro fertilization and re-derivation rate of non-treated embryos.

| Strain | # of examined females<sup>A</sup> | # of superovulated females<sup>B</sup> (A/B) | # of oocytes<sup>C</sup> | # of zygotes<sup>D</sup> (D/C) | # of cultured zygotes<sup>E</sup> | # of 2-cell stage embryos<sup>F</sup> (F/E) | # of transferred 2-cell embryos<sup>G</sup> | # of recipients<sup>H</sup> | # of pregnant recipients<sup>I</sup> (I/H) | # of newborns<sup>J</sup> (J/G) | # of weaned pups<sup>K</sup> (K/G) |
| --- | --- | --- | --- | --- | --- | --- | --- | --- | --- | --- | --- |
| BALB/c | 27 | 24 (88.9%) | 321 | 310 (96.6%) | 75 | 74 (98.7%) | 40 | 2 | 2 (100%) | 10 (25.0%) | 10 (25.0%) |
| NC | 20 | 19 (95.0%) | 328 | 319 (97.3%) | 43 | 43 (100%) | 40 | 2 | 2 (100%) | 28 (70.0%) | 26 (65.0%) |
| CBA | 27 | 23 (85.2%) | 239 | 222 (92.9%) | 153 | 150 (98.0%) | 40 | 2 | 2 (100%) | 18 (45.0%) | 18 (45.0%) |
| C3H | 12 | 12 (100%) | 261 | 256 (98.1%) | 86 | 85 (98.8%) | 40 | 2 | 2 (100%) | 11 (27.5%) | 11 (27.5%) |
| SJL | 27 | 26 (96.3%) | 503 | 479 (95.2%) | 187 | 182 (97.3%) | 40 | 2 | 2 (100%) | 20 (50.0%) | 18 (45.0%) |
| DBA1 | 24 | 23 (95.8%) | 510 | 428 (83.9%) | 308 | 307 (99.7%) | 40 | 2 | 2 (100%) | 14 (35.0%) | 14 (35.0%) |
| DBA2 | 20 | 16 (80.0%) | 432 | 415 (96.1%) | 75 | 74 (98.7%) | 60 | 3 | 3 (100%) | 10 (16.7%) | 10 (16.7%) |
| B6N | 32 | 29 (90.6%) | 502 | 473 (94.2%) | 207 | 206 (99.5%) | 60 | 3 | 2 (66.7%) | 18 (30.0%) | 18 (30.0%) |

### Zygote genome editing by electroporation in multiple inbred mouse strains

We chose *Hr* as our target gene because *Hr* deficient mice exhibit a hairless phenotype but show no abnormalities in prenatal development or postnatal growth (Benavides et al., 2009). We employed an exon deletion strategy to remove a critical region consisting of exon(s) shared by all annotated full-length transcripts to induce a frameshift mutation (Skarnes et al., 2011). Another strategy to induce a knockout is to target the coding sequence and introduce indel mutations; however, this approach can induce incomplete knockout mutations due to illegitimate translation (Makino et al., 2016; Hoshino et al., 2017).

We designed guide RNAs to excise exon3 of *Hr* (Fig. 1). Parallel experiment from IVF to the delivery of newborns using B6N zygotes were conducted with other 7 strains to confirm that no toxic effect of in vitro fertilisation, ribonucleoprotein (RNP) electroporation, or overnight culture was detected in B6N (Table S1). We found that except for CBA, all zygotes survived electroporation (97.1%, Table 2), exhibiting 2-cell development rates ranging from 47.1% (SJL) to 94.0% (CBA). Zygote electroporation resulted in decrease in the 2-cell rate in BALB/c (from 98.7% to 63.0%) as well as SJL (from 97.3% to 47.1%) (Tables 1 and 2). In addition, we observed a 100% pregnancy rate for all strains, except C3H (66.7%). Interestingly, we determined that birth rates were comparable to those achieved without electroporation in 5 out of 8 strains (BALB/c, C3H, SJL, DBA2, and B6N), whereas were clearly decreased in NC (from 70.0 % to 35.0%), CBA (from 45.0% to 28.3%), and DBA1 (from 35.0% to 20.0%) (Tables 1 and 2). G0 pups showed a hairless phenotype at approximately 3 weeks of age (Fig. 2), reflecting an abnormal second hair cycle (Zarach et al., 2004). We obtained pups with hair loss for all strains, with an incidence ranging from 42.9% (BALB/c) to 77.8% (B6N, Table 2).

**Fig. 1.**
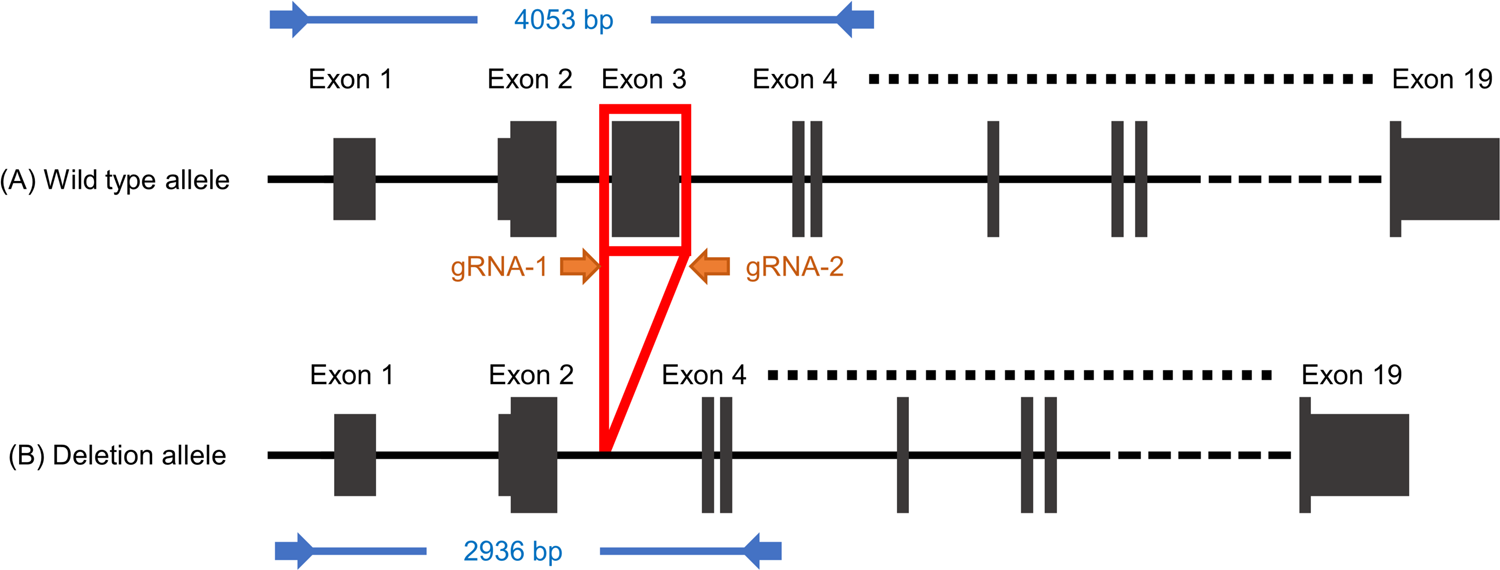
Genome editing and genotyping design on *Hr* gene. Orange arrows (gRNA-1 and gRNA-2) represent the gRNA target sites flanking exon 3. Blue arrows represent PCR primers, including the size of the PCR amplicon for G0 genotyping. Allele (A) represents the wild-type allele. whereas allele (B) represents the exon deletion allele.

**Fig. 2.**
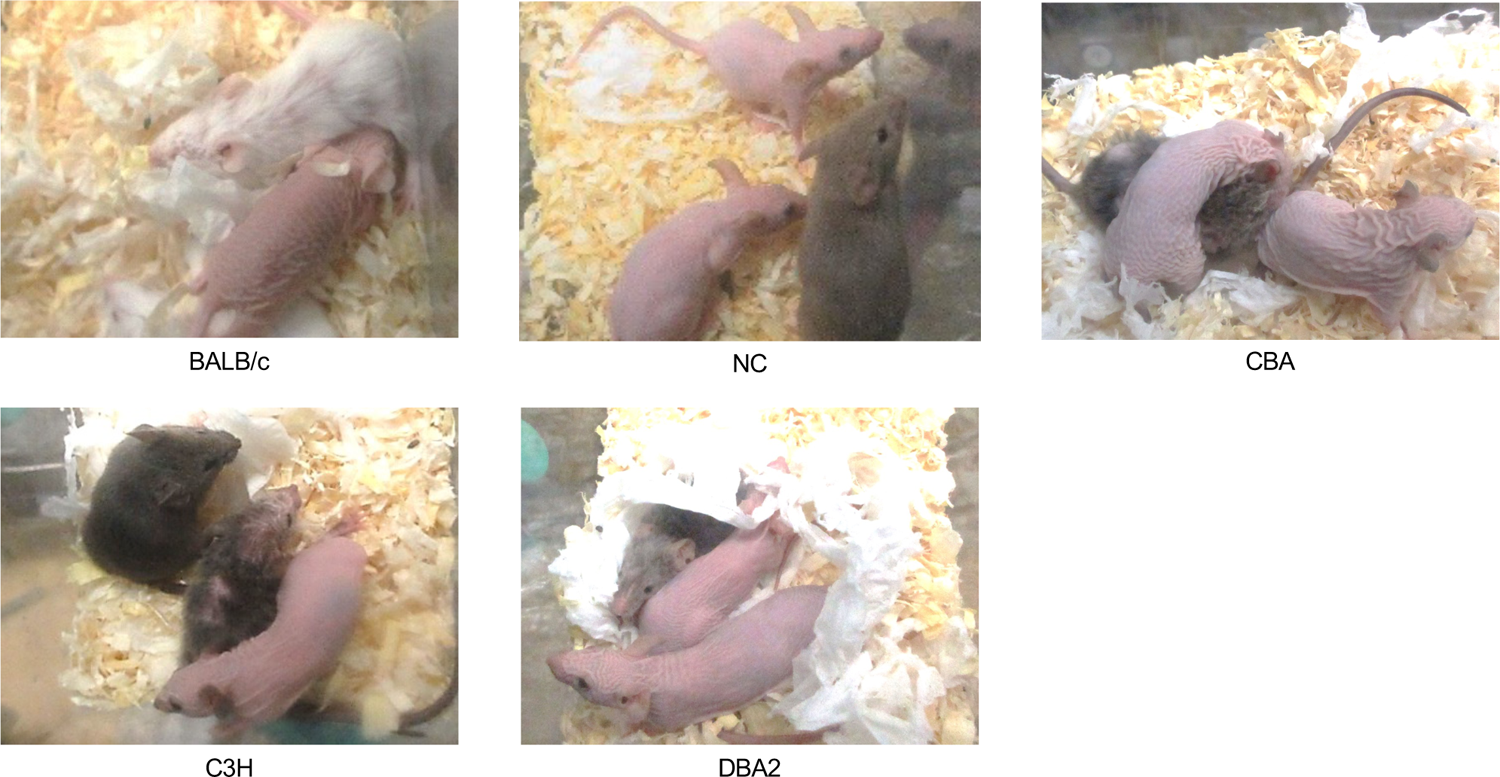
Phenotype of G0 mice in various inbred strains. Mice with induced mutations in the *Hr* gene begin to lose hair at approximately 3 weeks of age, when the second hair cycle begins. Complete loss-of-function of the *Hr* gene results in mice with complete hair loss.

**Table 2.**
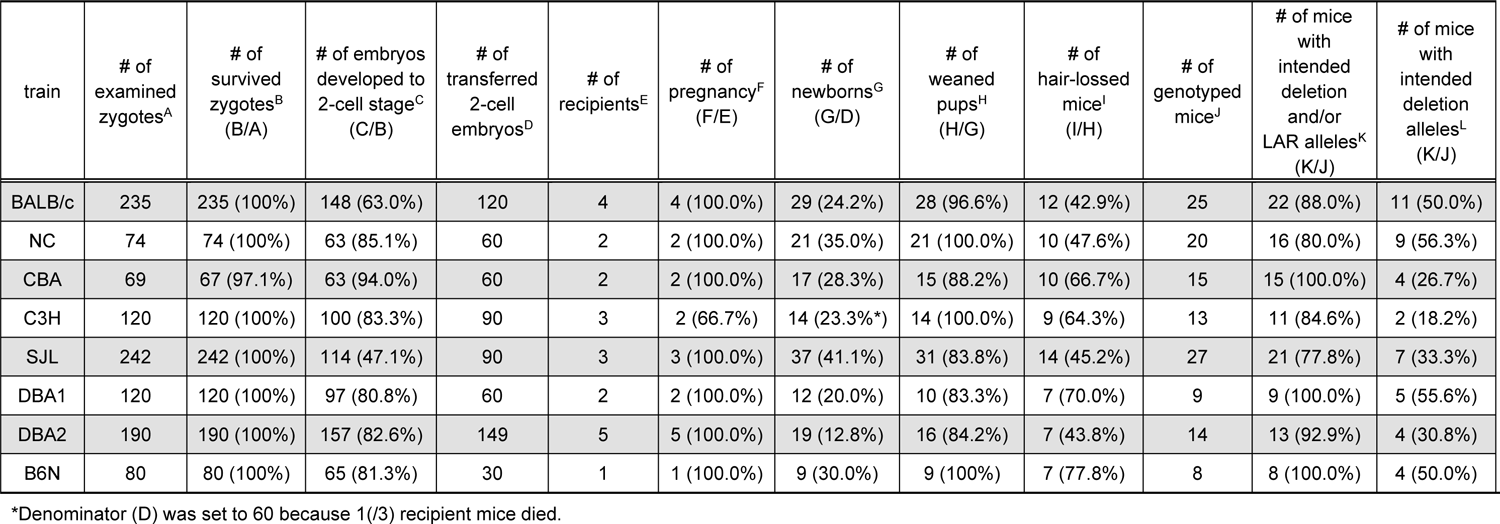
Survival rate, re-derivation rate, and editing efficiency of electroporated zygotes.

| train | # of examined zygotes<sup>A</sup> | # of survived zygotes<sup>B</sup> (B/A) | # of embryos developed to 2-cell stage<sup>C</sup> (C/B) | # of transferred 2-cell embryos<sup>D</sup> | # of recipients<sup>E</sup> | # of pregnancy<sup>F</sup> (F/E) | # of newborns<sup>G</sup> (G/D) | # of weaned pups<sup>H</sup> (H/G) | # of hair-lossed mice<sup>I</sup> (I/H) | # of genotyped mice<sup>J</sup> | # of mice with intended deletion and/or LAR alleles<sup>K</sup> (K/J) | # of mice with intended deletion alleles<sup>L</sup> (K/J) |
| --- | --- | --- | --- | --- | --- | --- | --- | --- | --- | --- | --- | --- |
| BALB/c | 235 | 235 (100%) | 148 (63.0%) | 120 | 4 | 4 (100.0%) | 29 (24.2%) | 28 (96.6%) | 12 (42.9%) | 25 | 22 (88.0%) | 11 (50.0%) |
| NC | 74 | 74 (100%) | 63 (85.1%) | 60 | 2 | 2 (100.0%) | 21 (35.0%) | 21 (100.0%) | 10 (47.6%) | 20 | 16 (80.0%) | 9 (56.3%) |
| CBA | 69 | 67 (97.1%) | 63 (94.0%) | 60 | 2 | 2 (100.0%) | 17 (28.3%) | 15 (88.2%) | 10 (66.7%) | 15 | 15 (100.0%) | 4 (26.7%) |
| C3H | 120 | 120 (100%) | 100 (83.3%) | 90 | 3 | 2 (66.7%) | 14 (23.3%*) | 14 (100.0%) | 9 (64.3%) | 13 | 11 (84.6%) | 2 (18.2%) |
| SJL | 242 | 242 (100%) | 114 (47.1%) | 90 | 3 | 3 (100.0%) | 37 (41.1%) | 31 (83.8%) | 14 (45.2%) | 27 | 21 (77.8%) | 7 (33.3%) |
| DBA1 | 120 | 120 (100%) | 97 (80.8%) | 60 | 2 | 2 (100.0%) | 12 (20.0%) | 10 (83.3%) | 7 (70.0%) | 9 | 9 (100.0%) | 5 (55.6%) |
| DBA2 | 190 | 190 (100%) | 157 (82.6%) | 149 | 5 | 5 (100.0%) | 19 (12.8%) | 16 (84.2%) | 7 (43.8%) | 14 | 13 (92.9%) | 4 (30.8%) |
| B6N | 80 | 80 (100%) | 65 (81.3%) | 30 | 1 | 1 (100.0%) | 9 (30.0%) | 9 (100%) | 7 (77.8%) | 8 | 8 (100.0%) | 4 (50.0%) |

\*Denominator (D) was set to 60 because 1(/3) recipient mice died.

### Genotyping of founder mice by nanopore long-read sequencing

Simultaneous two-site cleavage using Cas9 and a pair of guide RNA induces not only the intended regional deletion but also unexpected larger deletions (large rearrangements; LARs) and inversions (Canver et al., 2014). To precisely evaluate the genome-editing mutations carried in the G0 mice generated in the above experiment, we first performed long-PCR on 131 G0 mice of all 8 inbred strains. We confirmed the length of these PCR amplicons using agarose gel electrophoresis (Fig. S1). Individuals exhibiting at least a single band shorter than that of the wild-type were classified as mice with the intended deletion or LAR alleles (BALB/c, n = 22/25; NC, n =16/20; CBA, n = 15/15; C3H, n = 11/13; SJL, n = 21/27; DBA1, n = 9/9; DBA2, n = 13/14; B6N, n = 8/8; Table 2).

However, this evaluation cannot determine whether PCR bands reflect alleles with the intended deletion or unintended aberrant mutations. Therefore, we performed nanopore long-read sequencing and analysis using our software DAJIN (Kuno et al., 2022) to evaluate the presence of the intended alleles in each sample. We led DAJIN define “LAR” as a mutation with a deletion or insertion of 51 bp or more than the assumed intended deletion or wild-type sequence. Mutations within ±50 bp of the assumed intended deletion sequence were defined as “intended deletion” and mutations within ±50 bp of the wild-type sequence as “potential wild-type (pWT)”. We evaluated 116 G0 mice from all 8 inbred strains (BALB/c, n = 22; NC, n = 16; CBA, n = 15; C3H, n = 11; SJL, n = 21; DBA1, n = 9; DBA2, n = 13; and B6N, n = 8). DAJIN analysis revealed that 18.2% (C3H, n = 2/11) to 56.3% (NC, n = 9/16) of samples in each strain contained more than 10% of the intended deletion alleles (Fig. 3A and Table 2). Interestingly, we observed that NC mice were prone to have an intended deletion allele, whereas C3H mice tended to have LAR alleles (Fig. 3B and S2). These results demonstrated that our method induced the intended genome-editing mutations in all 8 inbred strains.

**Fig. 3.**
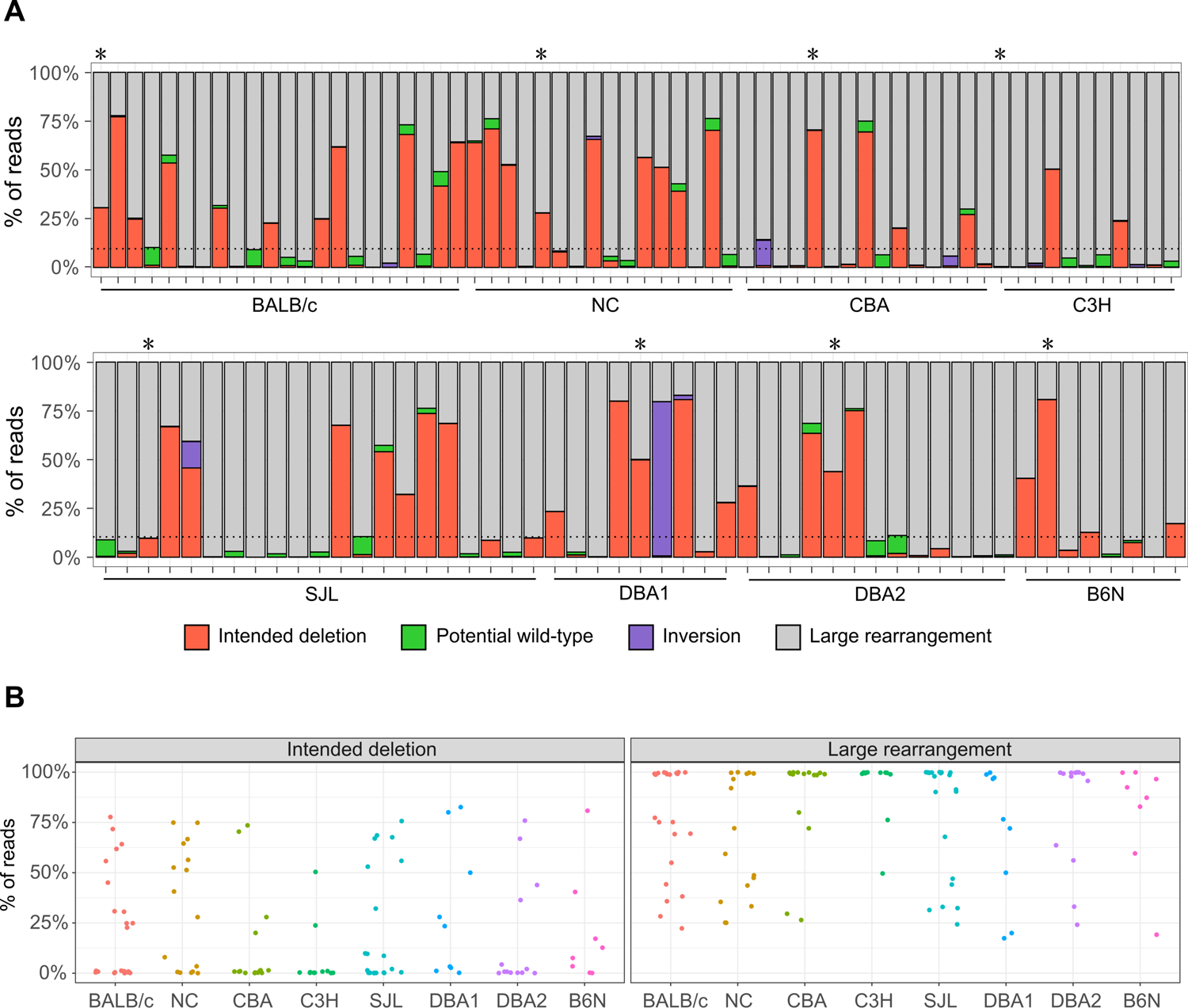
DAJIN report of the allele percentage. A. DAJIN report of the allele percentage in the 8 strains. The x-axis represents each mouse strain. The y-axis represents the percentage of reads with DAJIN predicted allele types. The bar colour represents each of DAJIN predicted allele types, including intended deletion, potential wild-type, inversion, and large rearrangement. Horizontal dots represent the 10% line of allele percentage. Asterisks denote G0 mice used in the creation of G1 mice. B. Scatter plots of the percentage of reads on the intended deletion and large rearrangement alleles. Dots represent each mouse sample. The x-axis and colours of dots represent mouse strains. The y-axis represents the percentage of reads with DAJIN predicted allele types.

### Confirmation of heritability of mutant alleles

Establishment of novel strains of mice is critical for in vivo experiments. As genome editing using mouse zygotes can often induce mosaicism (Mizuno et al., 2014), we cannot guarantee that the mutations detected in G0 mice are inherited by the next generation. To assess the differences in the establishment of mutant lines among inbred mouse strains using our method, we investigated the heritability of the mutant alleles carried by G0 mice. We thus used frozen sperm from 1 G0 male mouse of each inbred strain (Fig. 3A) for IVF. We confirmed the presence of PCR products that appeared to be derived from the intended deletion allele in all strains, except for C3H, in which the G0 mouse did not carry the intended deletion allele (Fig. 3A). Targeted amplicon short-read sequencing revealed that only a single type of the intended deletion allele (allele 1) was detected in BALB/c, NC, CBA, SJL, DBA2, and B6N mice, whereas 2 types of deletion alleles (alleles 1 and 2) were found in DBA1 mice (Fig. 4A and Table S2). The short-read sequencing also discovered the presence of a single type of the LAR allele (allele 2) in DBA2 mice. We determined that these intended deletion and LAR sequences matched perfectly with the sequence output from DAJIN analysis in each G0 mouse (Fig. 4B).

**Fig. 4.**
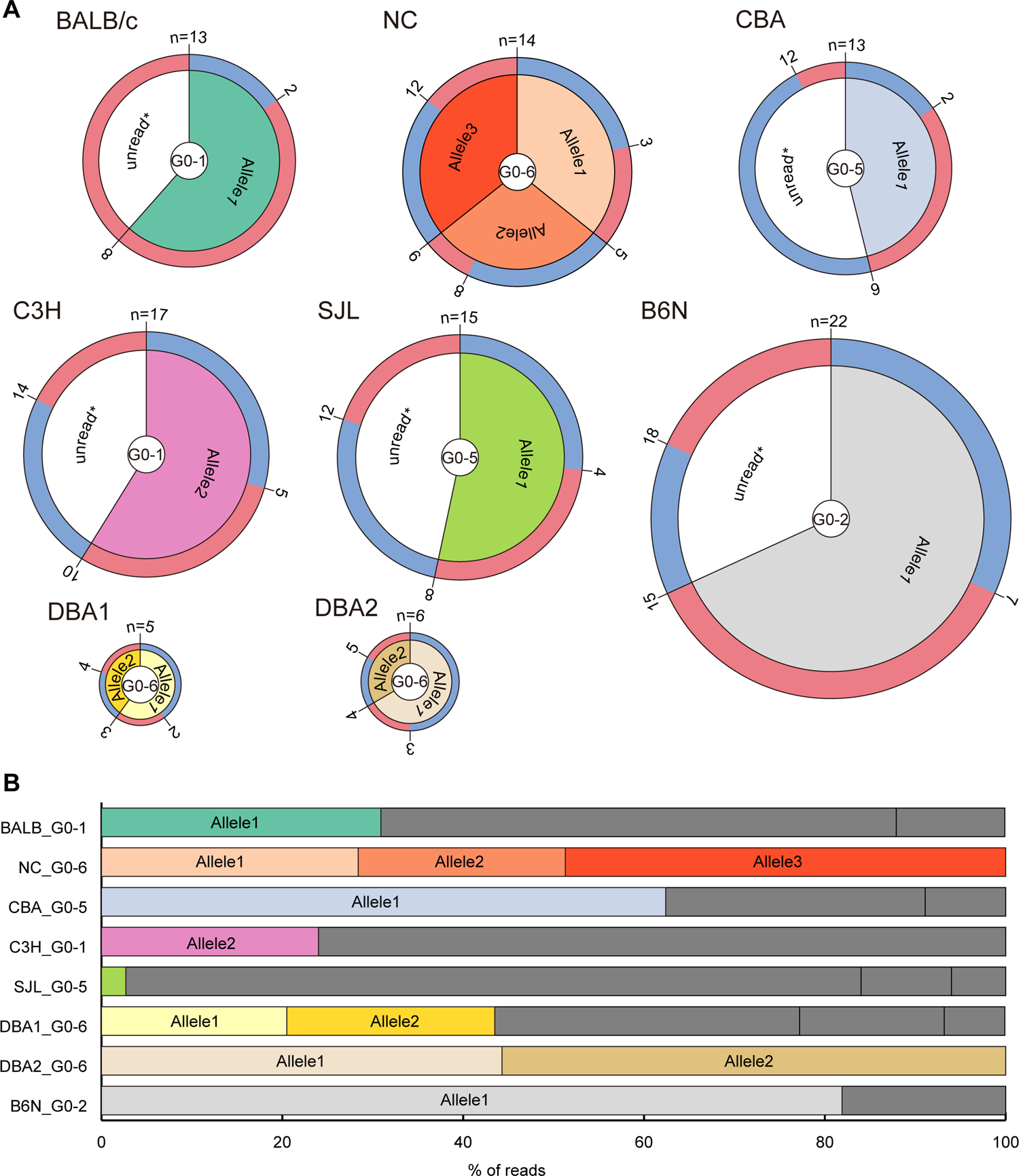
Heritability of genome editing mutations to progenies. A. The number, genotype, and sex of G1 mice of 8 inbred strains are shown. The outer layer represents sex (blue for male and red for female), the middle layer represents the allele carried by each G1 mouse, and the inner layer shows G0 male mice used for each strain. Unread* alleles might include potential wild-type, inversion, or large rearrangement alleles (not confirmed). The presence of these alleles was confirmed by electrophoresis of genomic PCR products. Note that all biological mothers of G1 mice were wild-type. B. Read percentages of mutant alleles harboured by each father of G1 mice based on DAJIN analysis. The bar colour links to the colour of each allele in G1 mice in Fig. 4A. Dark grey indicates unread* alleles that were not selected for detailed analysis in the genotyping of G1 mice.

As targeted amplicon short-read sequencing has a limited range of analysis, we used Sanger sequencing to check whether the LAR alleles in the G0 mice in C3H and NC strains were inherited by G1 mice. We accordingly detected 1 type (allele 2) in C3H mice and 2 types of LAR alleles (alleles 2 and 3) in NC mice, the sequences of which were in perfect alignment with the respective sequence output from DAJIN analysis in G0 mice (Fig. 4B). These results clearly showed that DAJIN outputs the exact sequence of each mutant allele in the G0 generation, further suggesting the establishment of mutant mouse lines using our method.

## Discussion

We report a universal zygote genome editing method capable of inducing critical exon elimination knockout in 8 inbred mouse strains. Although IVF efficiency and embryonic development rates were lower in a few strains, editing efficiency was sufficient to generate G0 mice and establish knockout strains. We detected both the intended deletion and unintended mutations (LAR and inversion) in our strains. We also confirmed that the mutant alleles detected in G0 mice were transmitted to the next generation. These results clearly demonstrated that our approach is suitable for generating mutant strains from the genetic backgrounds tested here.

The percentage of super-ovulated females, fertilisation rates, and embryo development rates in IVF and embryo transfer (Table 1) were not significantly different from those in previous reports (Byers et al., 2006). We did not examine the optimal conditions for performing super-ovulation treatments in each inbred strain, so there might be even better conditions for strains, such as BALB/c, which tend to have low fertilisation rates in a medium with low osmolarity or calcium concentration (Golkar-Narenji et al., 2012). The ovulation rate and number of oocytes obtained per female might be improved by hyper-ovulation treatment with anti-inhibin serum or in combination with oestrous cycle synchronisation (Hasegawa et al., 2022). We should emphasise that the embryo development rates at the 2-cell stage in BALB/c, C3H, and SJL mice (Table 1) were superior to those described in a previous study (Byers et al., 2006), implying that the IVF method using methyl-β-cyclodextrin and reduced glutathione enhanced fertility in these strains.

In addition, the survival sensitivity or embryonic development to electroporation stimuli differed among strains (Table 2). The development rate of 2-cell stage embryos was lower in the genome-edited group than in the non-genome-edited group in BALB/c (99% to 63%) and SJL (97% to 47%) mice compared with that in other strains. Moreover, the birth rates were lower in the genome-edited group than in the non-genome-edited group in NC (70% to 35%), CBA (45% to 28%), and DBA1 (35% to 20%). Compatibility to IVF conditions might also have an impact on survival rate and editing efficiency, as reduced glutathione has been suggested to weaken the zona pellucida and allow delivery of ribonucleoprotein components by electroporation (Wang et al., 2016). Further experiments targeting other loci, as well as comparative analysis between electroporation and microinjection, need to be conducted to reveal the differences of strains in sensitivity to electroporation.

For genotyping of G0 mice, long-read sequencing was used to analyse the broad genomic regions of a large number of mice. Although this technology is comprehensive, its weakness is the high rate of sequencing errors. We overcame this bottleneck by adopting our novel software DAJIN, which can automatically identify and classify both intended and unintended diverse mutations (Kuno et al., 2022). Mutant alleles in G0 samples detected by long-read sequencing and DAJIN were also detected in G1 mice. Long-read sequencing analysis suggested that the frequency of the appearance of the mutation pattern might be different in each lineage (Fig. 3B and S2). Although differences in mutation frequency need to be analysed at more target loci, this method has the potential to reveal the molecular (genetic background) and environmental (developmental speed) causes of toxic LARs.

The ability to generate knockout mice using a universal method will contribute to not only the establishment of a rapid, parallel bioresource, but also comparative phenotype analysis of knockout alleles among diverse inbred strains. Detailed phenotyping analysis of each *Hr* knockout strain established in our study will reveal strain difference in the *Hr* gene function. All 8 strains in this study are derived primarily from *Mus musculus domesticus* and are therefore genetically close to each other. It is hence important to also examine the feasibility of our method in *Mus musculus musculus*, *Mus musculus castaneus*, and *Mus spretus* (Low et al., 2022). Whether this method is useful for heterogeneous populations, such as the Collaborative Cross and Diversity Outbred (Churchill et al., 2004; Rasmussen et al., 2014) is also an interesting question. Our method can provide a series of mutant mouse lines from various genetic backgrounds with minimal artefacts such as off-target mutagenesis through genome editing, expanding the diversity of mammalian reverse genetics.

## Materials and Methods

### Animals

One outbred (Crl:CD1(ICR)) and 8 inbred (BALB/cAnNCrlCrlj, NC/NgaTndCrlj, CBA/J, C3H/HeNCrl, SJL/J, DBA/1JNCrlj, DBA/2NCrl, and C57BL/6NCrl) strains of mice (*Mus musculus*) were provided by the Jackson Laboratory Japan, Inc. (Yokohama, Japan). All mice were maintained under specific-pathogen-free (SPF) conditions, provided with water and steam-sterilised CRF-1 (Oriental Yeast, Tokyo, Japan) or irradiated CE-2 (CLEA Japan, Tokyo, Japan) ad libitum, and housed under controlled lighting conditions (daily light period, 06:00– 18:00 or 08:00–20:00). We observed the animals daily, and performed bedding and water changes once a week. On the day of IVF, the animals were euthanised by cervical dislocation and used in the experiments. At the end of maintaining colonies, all animals were euthanised by CO_2_ inhalation. All animal experiments were approved by the Institutional Animal Care and Use Committee (IACUC) at The Jackson Laboratory Japan, Inc. and the RIKEN Tsukuba Branch. *Hr* knockout mouse strains in each inbred background were deposited at RIKEN BioResource Research Center (BRC): RBRC11694 (BALB/c), 11697 (NC), 11696 (CBA), 11700 (C3H), 11695 (SJL), 11698 (DBA1), 11699 (DBA2), and 10692 (B6N).

### Super-ovulation

Female mice at 9 to 11 weeks of age or those from retired breeders were injected intraperitoneally with 7.5 IU pregnant mare serum gonadotropin (PMSG; PmsA, Nippon Zenyaku Kogyo Co., Ltd., Fukushima, Japan) followed by an injection of 7.5 IU human chorionic gonadotropin (hCG; Gonadotropin, ASKA Animal Health Co., Ltd., Tokyo, Japan) 48 h later. Between 16 to 18 h after hCG injection, mature metaphase II (MII) oocytes were collected from the ampulla region of the oviducts using a needle under paraffin oil and transferred to the fertilisation medium.

### In vitro fertilization

For the IVF experiments, we used FERTIUP sperm pre-culture medium containing methyl-β-cyclodextrin for sperm pre-culture and CARD MEDIUM containing reduced glutathione for fertilisation (Kyudo, Saga, Japan). Spermatozoa from the epididymal caudae of male mice (at 9 to 11 weeks of age or those from retired breeders) of each strain were suspended in 100 μL FERTIUP sperm pre-culture medium (Kyudo) and incubated at 37°C in an atmosphere of 5% CO_2_ for 30–60 min. Sperm from G0 mice were cryo-preserved in plastic straws using FERTIUP Sperm Cryopreservation Medium (Kyudo). Spermatozoa of G0 males were thawed and pre-incubated as previously described until used for IVF (Takeo and Nakagata, 2010). Collected cumulus-oocyte complexes (COCs) from female oviducts were pre-incubated in CARD MEDIUM for between 60 to 90 min. Subsequently, 2–6 μL of pre-incubated spermatozoa were transferred into 200 μL drops of CARD MEDIUM containing COCs, followed by incubation for insemination. Between 4 to 5 h after insemination, spermatozoa and cumulus cells were removed from the oocytes by pipetting. After washing twice with fresh medium, oocytes were cultured in KSOM medium (ARK Resource Ltd., Kumamoto, Japan) at 37°C and 5% CO_2_. Between 6 to 8 h after insemination, oocytes with 2 pronuclei were judged to be fertilised and subjected to genome editing experiments. Zygotes that were not used in embryo transfer or genome editing experiments were cryo-preserved by a simple vitrification method using DAP213 (ARK Resource).

### Embryo Transfer

Thirteen to fifteen embryos developed to the 2-cell stage after genome editing or 10 non-genome-edited 2-cells or pronuclear-stage zygotes from IVF using frozen-thawed sperm were transferred into each oviduct of day-1 pseudopregnant Crl:CD1(ICR) recipients. All embryo transfer experiments were performed under appropriate balanced anaesthesia with butorphanol (Vetorphale, Meiji Animal Health Co., Ltd., Kumamoto, Japan), medetomidine (Domitor, Nippon Zenyaku Kogyo Co., Ltd., Fukushima, Japan), and midazolam (MIDAZOLAM SANDOZ, Sandoz, Tokyo, Japan). Post-operative pain management was performed by subcutaneous administration of carprofen (Rimadyl; Zoetis Inc., Tokyo, Japan).

### Health and phenotype assessment

Heath assessment of pups was performed in their home cages at the age of weaning (3 to 4 weeks after birth) to detect any unexpected appearance and behaviours or dysmorphological characteristics. Phenotype analysis of hair loss in G0 mice was conducted between 3 to 6.5 weeks after birth prior to sequencing analysis. Investigators could not be blinded to the mouse strain due to coat color difference. Photographs were obtained between the age of 4 to 7 weeks by using IXY 600F or 650 digital cameras (Canon, Tokyo, Japan).

### Genome editing by electroporation

Cas9 protein, crRNAs, and tracrRNAs were purchased from Integrated DNA Technologies (Coralville, IA, USA). Two crRNAs were designed to target the *Hr* gene in 8 inbred strains (5′-CTA ACA CTT GGC ATG ACC AA-3′ and 5′-GAT GGA AGC CCC TGG CTA GA-3′). The RNP complex was prepared in Opti-MEM (Thermo Fisher Scientific, Waltham, MA, USA) with 2.4 μM Cas9 protein and each of 3.7 μM crRNA and 7.4 μM tracrRNA at 25°C for 5 min. RNP solutions were prepared immediately before electroporation and kept on ice until use.

Electroporation was performed using the TAKE method (Kaneko, 2017) with a NEPA21 Super Electroporator (NEPA GENE Co. Ltd, Chiba, Japan). The poring pulse was set to a voltage of 40 V, pulse length of 3.0 ms, pulse interval of 50 ms, number of 4 pulses, decay rate of 10%, and + polarity. The transfer pulse was set to a voltage of 10 V, pulse length of 50 ms, pulse interval of 50 ms, number of 5 pulses, decay rate of 40%, and +/− polarity. A 1-mm gap electrode (CUY501P1-1.5, NEPA GENE) was filled with 5 μL RNP solution, and zygotes washed with Opti-MEM solution were arranged on the electrode. Electroporated zygotes were observed for survival and cultured in KSOM overnight at 37°C and 5% CO_2_.

### Genomic DNA extraction

Ear clips of G0 mice were collected between 3 to 8 weeks after birth. Genomic DNA was extracted using Lyppo (Gene modification, Osaka, Japan) or DNeasy Blood & Tissue Kits (Qiagen, Venlo, Netherlands) according to the manufacturer’s protocols (Table S3).

Genomic DNA was extracted from G1 mouse tail tips using the conventional phenol/chloroform/isoamyl alcohol (PCI) method. Briefly, samples treated with proteinase K solution (NACALAI TESQUE, Inc., Kyoto, Japan) were purified by PCI extraction, followed by ethanol precipitation, and resuspended in TE buffer solution (pH 8.0) (NACALAI TESQUE).

### Long-PCR and long-read sequencing

Long PCR amplification of on-target genomic DNA regions was performed using purified genomic DNA, KOD multi&Epi (TOYOBO Co., Ltd., Osaka, Japan), and appropriate primers (Table S4). The products were loaded into 1.5% agarose gel (NIPPON GENE, Tokyo, Japan) containing ethidium bromide (Thermo Fisher Scientific) at final concentration of 0.4 µg/ml. After electrophoresis under 100V for 40 to 45 min, PCR products were visualized using FAS-V imaging system (NIPPON Genetics, Tokyo, Japan).

Nested PCR for imparting barcode sequences was performed using a 5-fold dilution of the first PCR products with distilled water, KOD multi&Epi, and appropriate primers (Table S4). We designed 72 primer sets for DNA barcoding, of which 69 pairs were used. Equal amounts of barcoded nested PCR products were mixed and purified using the FastGene Gel/PCR Extraction Kit (NIPPON Genetics). Purified PCR products (30 ng/µL) were used to prepare the nanopore long-read sequencing library. Library preparation was performed using the NEBNext End repair/dA-tailing Module (New England BioLabs, Ipswich, MA, USA) and the Ligation Sequencing 1D kit SQK-LSK109 (Oxford Nanopore Technologies, Oxford, UK) according to the manufacturer’s protocol. The prepared library was loaded into the R9.4 SpotON Flow Cell_FLO-MIN106 (Oxford Nanopore Technologies), and a MinKNOW GUI (version 22.03.06) sequence run was performed for 36 h.

Nanopore sequencing reads were base-called and de-multiplexed using Guppy version 6.1.3+cc1d765d3 (Oxford Nanopore Technologies). Alleles in each sample were classified using DAJIN version 0.6.0 (Kuno et al., 2022). Stack and point plots of read percentages in each sample were produced using custom Bash and R scripts (https://github.com/akikuno/knockout-in-different-strains). Mice with more than 10% target deletion alleles were defined as target mice. For visualisation, nanopore reads were mapped against chromosome 14 of the mouse genome assembly GRCm38.p6 using minimap2 version 2.22-r1101 with options ‘-ax map-ont’ (Li, 2018). Mapped reads were visualised using IGV version 2.13.1 (Robinson et al., 2011).

### Short-PCR and short-read next-generation sequencing

Genomic short-PCR was performed using AmpliTaq Gold 360 DNA Polymerase (Thermo Fisher Scientific). The primers used are listed in Table S4. Nested PCR for adding the barcode sequence was performed using AmpliTaq Gold 360 DNA Polymerase and relevant primers whose barcode sequences were added to the 5′ end of targeted *Hr* mutated amplicons (Table S4). Nested PCR amplicons were purified using 1.12X AMPure XP beads (Beckman Coulter Genomics, Brea, CA, USA). Ten percent spike-in of PhiX control V3 (Illumina, San Diego, CA, USA) was added to these amplicons. Paired-end sequencing (2 × 150 bases) of these amplicons was performed using an iSeq 100 (Illumina).

Sequencing reads were de-multiplexed using the GenerateFASTQ module version 2.0.0 on iSeq 100 Software (Illumina). Analysis of on-target amplicon sequencing was performed using CRISPResso2 version 2.2.9, in batch mode (Clement et al., 2019).

### Sanger sequencing

PCR was performed using purified genomic DNA, AmpliTaq Gold 360 DNA polymerase, and appropriate primers (Table S4). PCR amplicons were purified using the FastGene Gel/PCR Extraction Kit (NIPPON Genetics). Sequencing reactions were performed using Purified DNA fragments, the BigDye Terminatorv3.1 Cycle sequencing Kit (Thermo Fisher Scientific), and appropriate primers (Table S4). A 3500 Genetic Analyser (Thermo Fisher Scientific) was used for Sanger sequencing analysis.

## Acknowledgements

We thank Mizuho Iwama, Masayo Kadota, Noriko Hiraiwa, and all other members of the Experimental Animal Division of RIKEN BRC and Yuta Fujiki of The Jackson Laboratory Japan, Inc. for their assistance.

## Data availability

All sequence data (FASTQ) (Nanopore and Illumina iSeq) from this study can be retrieved from SRA under the BioProject ID PRJNA874336 and PRJNA874320, respectively.

